# Loss of *Mfn1* but not *Mfn2* enhances adipogenesis

**DOI:** 10.1101/2022.11.04.515167

**Authors:** JP Mann, LC Tabara, A Alvarez-Guaita, L Dong, A Haider, K Lim, P Tandon, JEN Minchin, S O’Rahilly, S Patel, DJ Fazakerley, J Prudent, RK Semple, DB Savage

## Abstract

**Objective:** A biallelic missense mutation in mitofusin 2 (*MFN2*) causes multiple symmetric lipomatosis and partial lipodystrophy, implicating disruption of mitochondrial fusion or interaction with other organelles in adipocyte differentiation, growth and/or survival. In this study, we aimed to document the impact of loss of mitofusin 1 (*Mfn1*) or 2 (*Mfn2)* on adipogenesis in cultured cells.

**Methods:** We characterised adipocyte differentiation of wildtype (WT), *Mfn1*^*-/-*^ and *Mfn2*^*-/-*^ mouse embryonic fibroblasts (MEFs) and 3T3-L1 preadipocytes in which Mfn1 or 2 levels were reduced using siRNA.

**Results:** *Mfn1*^*-/-*^ MEFs displayed striking fragmentation of the mitochondrial network, with surprisingly enhanced propensity to differentiate into adipocytes, as assessed by lipid accumulation, expression of adipocyte markers (*Plin1, Fabp4, Glut4, Adipoq*), and insulin-stimulated glucose uptake. RNA sequencing revealed a corresponding pro-adipogenic transcriptional profile including *Pparg* upregulation. *Mfn2*^*-/-*^ MEFs also had a disrupted mitochondrial morphology, but in contrast to *Mfn1*^−/-^ MEFs they showed reduced expression of adipocyte markers and no increase in insulin-stimulated glucose uptake. *Mfn1* and *Mfn2* siRNA mediated knockdown studies in 3T3-L1 adipocytes generally replicated these findings.

**Conclusions:** Loss of *Mfn1* but not *Mfn2* in cultured pre-adipocyte models is pro-adipogenic. This suggests distinct, non-redundant roles for the two mitofusin orthologues in adipocyte differentiation.

## 1. Introduction

Primary mitochondrial disorders result in pleiotropic syndromes disturbing the function of multiple organ systems commonly including the central and peripheral nervous system and neuromuscular function[1,2]. Adipose tissue dysfunction and related features of the metabolic syndrome are not present in most primary mitochondrial disorders[3,4]. One striking exception to this is the recent discovery that a point mutation in human mitofusin 2 (*MFN2*) produces profound adipose tissue dysfunction, non-alcoholic fatty liver disease, insulin resistance and type 2 diabetes[5–7]. Humans with biallelic p.Arg707Trp MFN2 mutations develop increased upper body adiposity (multiple symmetric lipomatosis) with lower body lipodystrophy. Over-grown adipose depots show deranged ultrastructure of the mitochondrial network but largely normal adipocytes on light microscopy[5]. Affected adipose tissue exhibits a strong transcriptomic signature of activation of the integrated stress response and increased pro-growth signalling. This adipose phenotype appears to be specific to the p.Arg707Trp mutation, which has been identified in all reported cases to date. Many other *MFN2* mutations, many effectively leading to loss of one allele (i.e. haploinsufficiency) and often located in the N-terminal GTPase domain, are associated with autosomal dominant sensorimotor neuropathy without an adipose phenotype[8]. No monogenic disease attributable to *MFN1* mutations has yet been reported.

Within the last 20 years, the dynamic nature of the mitochondrial network has been clearly established, and many of the molecules mediating mitochondrial morphology have been identified[9,10]. Mitochondrial network dynamics are entrained to adapt to the cellular nutritional state: starvation or nutrient excess trigger changes in mitochondrial morphology that correspond to functional changes in mitochondrial oxidative phosphorylation[11] and the generation of reactive oxygen species (ROS)[12,13]. The mitofusins (*MFN1* and *MFN2*) are GTPases required for fusion of the outer mitochondrial membrane. Loss of either mitofusin leads to fragmentation of the mitochondrial network and disrupts its function[14–16]. *Mfn1* is widely expressed and is required for mitochondrial fusion whereas *Mfn2* reportedly has additional roles, such as being intimately involved in forming membrane contact sites between mitochondria and the endoplasmic reticulum[17] or lipid droplets[18], and in facilitating apoptosis and mitophagy[19].

Global *Mfn1* or *Mfn2* deletion in mice is embryonically lethal, but knockout of *Mfn2*[20–22] specifically in mature adipocytes has illustrated the importance of mitochondrial network dynamics in adipose tissue. Loss of *Mfn2* from *Ucp1*-expressing brown adipose tissue impairs its thermogenic capacity and increases lipid droplet size[20,21], but surprisingly increases insulin sensitivity. In contrast, deletion of *Mfn2* from mature white adipocytes in adult mice was obesogenic and impaired insulin sensitivity [22]. Adipose-specific knock-out of *Mfn1* has yet to be described. Importantly, all promoters used for adipose-specific knockouts to date have been from genes expressed during terminal adipocyte differentiation, so whether loss of one or both mitofusins perturbs the process of adipogenesis, which involves significant expansion of mitochondrial mass[23–25], is unknown.

Differentiation of fibroblast-like cells into mature adipocytes is a well-established experimental paradigm used to dissect molecular regulation of adipogenesis[26,27]. In this study we examined the role of *Mfn1* and *Mfn2* in adipogenesis using knock-out mouse embryonic fibroblasts (MEFs), validating findings in 3T3-L1 preadipocytes using siRNA-mediated knock-down. We found that loss of *Mfn1* enhances adipogenesis, whereas loss of *Mfn2* has no effect on total lipid accumulation but alters lipid droplet morphology and reduces the expression of some adipogenic markers. These findings suggest that the two mitofusin proteins have important but divergent roles in the control of adipogenesis.

## 2 Methods

### 2.1 Culture and adipogenic differentiation of MEFs

Mouse embryonic fibroblasts (MEFs) null for *Mfn1*^*-/-*^, *Mfn2*^*-/-*^, *Mfn1*^*-/-*^*2*^−/-^, and *Opa1*^*-/-*^, and wild-type MEFs, originally derived by Chen *et al*.[14], were obtained for use in this study (see Supplementary Table 1 for a list of reagents).

MEFs were cultured in Dulbecco’s Modified Eagle’s Medium supplemented with 10% (vol/vol) foetal bovine serum (FBS), 1% sodium pyruvate, 1% non-essential amino acids, 1% penicillin/streptomycin, 2mM L-glutamine, and 50mM 2-mercaptoethanol. All cells were grown at 37°C in 20% O_2_ and 5% CO_2_.

Cells were maintained and passaged whilst sub-confluent prior to adipogenic differentiation. To initiate differentiation, cells were grown to confluence (day -2) in 6- or 12-well plates and then, two days later, treated with 8μg/mL D-pantothenic acid, 8μg/mL biotin, 1μM rosiglitazone, 0.5mM 3-isobutyl-1-methylxanthine (IBMX), 1μM dexamethasone, and 1μM insulin. Treatment with D-pantothenic acid, biotin, rosiglitazone, and insulin was repeated on day +2, +4, and +6. Differentiation continued until day +8.

For fluorometric quantification of intracellular lipid accumulation, cells were incubated with AdipoRed at 1:25 dilution and 37°C for 20 minutes in the dark. Cells were washed twice in PBS then fluorescence was measured on Tecan M1000 Pro plate reader with excitation at 485nm and emission at 572nm.

### 2.2 Over-expression of *Pparg2* in MEFs

To improve the adipogenic potential of MEFs, cells were retrovirally transduced with murine Peroxisome proliferator-activated receptor gamma 2 (*Pparg2*). Phoenix-AMPHO (ATCC) cells were used for packaging. 70% confluent Phoenix cells were transfected with 12μg pBABE (puromycin-resistant)-mPparg2 vector or pBABE-EGFP (control) plasmid DNA using Lipofectamine 3000 transfection reagent. 72-hours later the supernatant media containing the virus was collected and filtered through a 0.45μm membrane. Retroviral stocks were used to transduce 50-60% confluent MEFs with addition of 12μg/mL polybrene. From 24-hours after transduction, puromycin selection began using 4μg/mL, and MEFs were subsequently cultured in media containing this concentration of puromycin. Adipogenic differentiation was then performed according to the above protocol.

### 2.3 Confocal microscopy of mitochondrial network

Cells were stained with Mitotracker Orange CMTMRos (ThermoFisher) for imaging of the mitochondrial network using confocal microscopy. MEFs were grown on cover slips to 80-90% confluence (pre-adipocytes at day -2 for MEFs and day 0 for 3T3-L1s) and treated with 400nM Mitotracker for 20 minutes at 37°C in the dark. Cells were washed with fresh media four times, each for 30 minutes at 37°C in the dark, followed by three washes in phosphate buffered saline (PBS). Cells were then fixed in 4% formaldehyde at room temperature for 10 minutes and mounted onto slides using ProLong Gold or Diamond Antifade Mountant with 4′,6-diamidino-2- phenylindole (DAPI) (ThermoFisher). Images were obtained using a Leica SP8 confocal microscope using excitation/emission spectra: Mitotracker Orange CMTMRos (550/560-580), BODIPY (490/515-535), and DAPI (405/450-470).

### 2.4 Culture and adipogenic differentiation of 3T3-L1 fibroblasts

Mouse fibroblast 3T3-L1 fibroblasts (3T3-L1s)[28] were used as an in vitro model of adipocyte differentiation and were obtained as a gift from D.J. Fazakerley and D.E. James (University of Sydney), originally obtained from Dr. Howard Green (Harvard Medical School, Boston, MA). 3T3-L1s were cultured in Dulbecco’s Modified Eagle’s Medium supplemented with 10% (vol/vol) foetal bovine serum and 1% GlutaMax. Cells were grown at 37°C in 20% O_2_ and 10% CO_2_. Cells were maintained and passaged sub-confluent prior to adipogenic differentiation.

To initiate differentiation, cells were grown to confluence (day -2) in 12-well plates and then two days later (day 0) treated with 0.5mM IBMX, 410nM biotin, 220nM dexamethasone, and 350nM insulin. After 72-hours (day +3), media was replaced with addition of 350nM insulin. A further 72-hours later (day +6) media was changed (without any additional chemicals) and subsequently replaced on day +8 and day +10 with experiments ending on day +12. On day +12, lipid accumulation was assessed using Oil Red O staining and AdipoRed quantification. RNA and protein were also extracted at day +12.

### 2.5 siRNA knock-down experiments in 3T3-L1s

For knock-down, confluent pre-adipocyte (day -2) 3T3-L1s grown in 12-well plates were transfected with a pool of two anti-Mfn1 siRNAs (ThermoFisher Scientific, CatIDs: s85002 & s85004), each at 40nM to achieve a final concentration of 80nM, with 2.5μl RNAiMax (ThermoFisher Scientific) in OptiMEM (ThermoFisher Scientific). For knock-down of Mfn2, an anti-Mfn2 SmartPool (Dharmacon) was used at a final concentration of 40nM. In all experiments, a scrambled Silencer Negative Control siRNA (ThermoFisher Scientific) was used at 80nM. Knock-down was performed with each media change, i.e. day -2, 0, +3, +6, +8, and +10.

### 2.6 Immunofluorescence in 3T3-L1s

Mature (day +8) differentiated 3T3-L1 adipocytes were reseeded onto matrigel-coated (Corning) coverslips at 50% density. siRNA knockdown was performed during reseeding. Reseeded adipocytes (day 9) were cultured under standard “high glucose” conditions (Dulbecco’s modified eagle’s medium containing 25 mM glucose supplemented with 10% (vol/vol) foetal bovine serum and 1% GlutaMax) or “low glucose” (serum-free Dulbecco’s modified eagle’s medium containing 5 mM glucose supplemented with 0.2% (w/v) bovine serum albumin and 1% GlutaMax) conditions for 16 h. Cells were then fixed in 4 % paraformaldehyde (PFA) in PBS at room temperature for 20 min, quenched with 100 mM glycine in PBS, washed 3 times with PBS, followed by incubation with 50 mM ammonium chloride in PBS to quench the unspecific fluorescence signal from aldehyde groups. Cells were washed again 3 times with PBS and permeabilized in 0.1 % Triton X-100 in PBS for 10 min. Then, cells were blocked with 10 % FBS in PBS, followed by incubation with the appropriate primary antibodies in 5 % FBS in PBS, for 2 hours at RT. After 3 washes with 5 % FBS in PBS, cells were incubated with specific secondary antibodies (1:1,000) for 1 hour at RT. After 3 washes in PBS, coverslips were mounted onto slides using Dako fluorescence mounting medium (Dako).

Mitochondria were stained using a rabbit anti-TOM20 antibody (11802-1-AP/Proteintech) (1:1,000). Alexa Fluor 488 (anti-rabbit) was used as a secondary antibody (1:1,000) (Invitrogen). Lipid droplets were stained by incubating coverslips with LipidTOX deep red (Thermo-fisher) for 2 hours (1:1,000).

For confocal image acquisition, cells were visualized using a 100X objective lenses (NA1.4) on a Nikon Eclipse TiE inverted microscope with appropriate lasers using an Andor Dragonfly 500 spinning disk system, equipped with a Zyla 4.2 PLUS sCMOS camera (Andor), coupled with Fusion software (Andor).

For image analysis, mitochondrial morphology was manually analysed and classified as intermediate, hyperfused or fragmented as already described[29]. The different mitochondrial parameters (area, length and number) were quantified by randomly selecting region of interests (ROIs) of 225 μm^2^ at the cell periphery and analysed using MitoMapr (Fiji)[30]. Error bars displayed on graphs represent the mean ± S.D from 3 independent experiments (at least 18 cells per experiment and condition). Statistical significance was analysed using Nested t-test or Two-way ANOVA test using GraphPad Prism software. *p < 0.05, **p < 0.01, *** p < 0.001 and **** p < 0.0001 were considered significant.

### 2.7 Electron microscopic imaging of mitochondria

Transmission electron microscope (TEM) studies were performed at the Cambridge Advanced Imaging Centre, University of Cambridge. Undifferentiated MEFs (day -2) were grown to sub-confluence in 6-well plates. Cells were washed twice in 0.9% NaCl then fixed in (2% glutaraldehyde/2% formaldehyde in 0.05M sodium cacodylate buffer pH 7.4 containing 2mM calcium chloride) for 4-hours at 4 °C. Cells were then scraped from plates and, after washing five times with 0.05 M sodium cacodylate buffer pH 7.4, samples were osmicated (1% osmium tetroxide, 1.5 % potassium ferricyanide, 0.05M sodium cacodylate buffer pH 7.4) for 3 days at 4°C. After washing five times in DIW (deionised water), samples were treated with 0.1% (w/v) thiocarbohydrazide/DIW for 20 minutes at room temperature in the dark. After washing five times in DIW, samples were osmicated a second time for 1-hour at room temperature (2% osmium tetroxide/DIW). After washing five times in DIW, samples were block stained with uranyl acetate (2% uranyl acetate in 0.05M maleate buffer pH 5.5) for 3 days at 4°C. Samples were washed five times in DIW and then dehydrated in a graded series of ethanol (50%/70%/95%/100%/100% dry) 100% dry acetone and 100% dry acetonitrile, three times in each for at least 5 minutes. Samples were infiltrated with a 50/50 mixture of 100% dry acetonitrile/Quetol resin (without BDMA) overnight, followed by 3 days in 100% Quetol (without BDMA). Then, samples were infiltrated for 5 days in 100% Quetol resin with BDMA, exchanging the resin each day. The Quetol resin mixture is: 12g Quetol 651, 15.7g NSA, 5.7g MNA and 0.5g BDMA (all from TAAB). Samples were placed in embedding moulds and cured at 60°C for 3 days.

Thin sections were cut using an ultramicrotome (Leica Ultracut E) and placed on bare 300 mesh copper TEM grids. Samples were imaged using a Tecnai G2 TEM (FEI/Thermo Fisher Scientific) run at 200 keV using a 20μm objective aperture to improve contrast. Images were acquired using an ORCA HR high resolution CCD camera (Advanced Microscopy Techniques Corp, Danvers USA).

Analysis was performed by manual measurement of individual mitochondria from all obtained images using ImageJ. Despite multiple mitochondria measured per sample, statistical tests (pairwise comparisons relative to wild-type) were based on the mean from two independent biological replicates and false-discovery rate adjustment for the number of tests[31].

### 2.8 mtDNA content assay

Relative mtDNA content was assayed using real-time quantitative polymerase chain reaction (RT-qPCR) for a mitochondrial DNA gene (*Rnr2*) and a nuclear DNA gene (*Hk2*). DNA was extracted from undifferentiated, MEFs at 80% confluence using DNeasy kit (Qiagen). DNA was quantified on a Nanodrop and diluted to 4ng/μl. RT-qPCR was performed in triplicate for each sample using 8ng DNA with primers for *Hk2* (Forward: GCCAGCCTCTCCTGATTTTAGTGT, Reverse: GGGAACACAAAAGACCTCTTCTGG) and *Rnr2* (Forward: AACTCGGCAAACAAGAACCC, Reverse: CCCTCGTTTAGCCGTTCATG). The ratio mt*Rnr2*/n*Hk2* was calculated using the standard curve method and expressed relative to wild-type.

### 2.9 Oil Red O staining

Mature adipocytes (day +8 MEFS or day +12 3T3-L1s) in six-well plates were gently washed in PBS before fixing in 10% formalin for 30 minutes at room temperature. Cells were then washed twice in PBS before dehydration in 60% isopropanol for 5 minutes (twice). Oil Red O (ORO) working solution was then added for 20 minutes. ORO working solution was made from 1g ORO (Sigma) dissolved in 400ml isopropanol, diluted 3:2 with milli-Q water and filtered through a 0.45μm membrane. Cells were washed with water then imaged on a flat-bed scanner and with light microscopy.

### 2.10 2-deoxy-glucose uptake assay

2-deoxy-glucose (2DOG) uptake assays were performed on mature adipocytes (day +8) following differentiation of MEFs in 24-well plates, according to standard protocols[32]. Briefly, MEFs were serum starved in Dulbecco’s Modified Eagle’s Medium (DMEM)/0.2% bovine serum albumin (BSA)/1% GlutaMax for 2-hours then washed three times in KRP buffer with 0.2% BSA. KRP buffer was prepared as: 0.6mM Na_2_HPO_4_, 0.4mM NaH_2_PO_4_, 120mM NaCl, 6 mM KCl, 1mM CaCl_2_, 1.2mM MgSO_4_ and 12.5mM HEPES adjusted to pH 7.4. At least one well on each plate served as a negative control by addition of 25μM cytochalasin B (Sigma Aldrich) to block all transporter-mediated glucose uptake. Cells were incubated with KRP/0.2% BSA ±100nM insulin for 20 minutes. After 15 minutes, 2-deoxyglucose (Sigma/Merck; radiolabelled 2DOG from PerkinElmer) was added to each well to a final concentration of 50 μM and 0.25 μCi/well. 5 minutes after addition of 2DOG, cells were quickly washed three times in ice-cold PBS then lysed in 1% (v/v) Triton X-100. Uptake of 2DOG (^3^H) was quantified using liquid scintillation counting and normalised to protein content.

### 2.11 RNAseq in knock-out MEFs

Bulk RNA sequencing (RNAseq) was performed on wild-type (WT) and Mfn1^−/-^ MEFs at four time points: day -2 (pre-adipocyte), day 0 (initiation of adipogenic differentiation), day +3 (early adipocyte), and day +8 (mature adipocyte). RNAseq was performed on Mfn2^−/-^ MEFs at two time points: day -2 (pre-adipocyte) and day +8 (mature adipocyte). Three independent biological replicates of each sample were used for analysis.

Cells were cultured and differentiated as described above, then briefly washed in PBS and lysed in RLT Buffer (Qiagen) with 1% 2-mercaptoethanol then passed through Qiashredder tubes (Qiagen). RNA was extracted using the RNeasy Isolation Kit (Qiagen) according to the manufacturers’ protocol. RNA was quantified using Agilent 2100 Bioanalyzer (Agilent Technologies Inc) and only samples with RNA Integrity Number ≥8 were used for library preparation. cDNA libraries were made using Illumina TruSeq RNA sample kits and sequencing was performed on Illumina NovaSeq 6000 with paired-end 150bp reads (Novogene, Cambridge, UK). Raw reads all passed quality control for Q_score_, error rate distribution, and AT/GC distribution.

Adapter sequences were removed from raw FASTQ files using cutadapt[33] and aligned to *Mus musculus* reference genome (GRCm38) using STAR[34]. Binary alignment/map (BAM) files were sorted using samtools[35] and counts were performed using featureCounts[36]. Differential gene expression (DGE) was performed using DESeq2[37], where significance was considered as a Benjamini-Hochberg false-discovery rate (FDR) corrected p-value <.01.

Pathway analysis was performed with the EnrichR package for R[38–40] using significantly differentially expressed genes to determine enriched Hallmark gene sets[41]. Gene sets with FDR-corrected p-value <.05 were considered enriched. Figures were generated in R 4.0.2[42] using packages pheatmap, ggplot2, and dplyr. RNAseq data is available in Supplementary Data 1. All code used in analysis is available from: https://doi.org/10.5281/zenodo.5770057.

### 2.12 Western blot studies

Cells were washed in cold PBS and lysed in RIPA buffer (Sigma) with PhosSTOP Phosphatase Inhibitor (Roche) and Complete-Mini Protease Inhibitor (Roche). Lysates were spun at 13,000rpm at for 15 minutes at 4°C then protein concentration quantified using BioRad DC Protein Assay Kit. 30-45μg protein lysates were mixed with NuPAGE 4x LDS buffer (ThermoFisher Scientific), containing 0.05% 2-mercaptoethanol, and denatured for 10 minutes at 95°C. Samples were run on 4-12% Bis-Tris gels (Invitrogen) and transferred onto a nitrocellulose membrane using iBlot-2 (ThermoFisher Scientific). Membranes were washed in Tris-buffered saline with 0.1% (vol/vol) Tween 20 (TBST, Sigma) before blocking in 5% (wt/vol) skimmed milk powder dissolved in TBST. Membranes were incubated with primary antibodies (Supplementary Table 2) at 4°C for 16-hours. Membranes were then washed with TBST five times for 5 minutes, followed by incubation with horseradish peroxidase (HRP)-conjugated secondary antibodies for 1-hour at room temperature. Blots were developed using Immobilon Western Chemiluminescent HRP Substrate (Millipore) with images acquired on BioRad ChemiDoc Imaging system.

### 2.13 Statistical analysis

Continuous data were expressed as mean ± standard deviation. Normally distributed data were analysed by t-test (for two group pairwise comparison) and one-way ANOVA (for three or more groups) with post-hoc Bonferroni multiple comparisons test, where FDR-corrected p-value <0.05 was considered significant. Details of specific analyses and the number of replicates performed are reported in figure legends. All experiments were conducted at least three times using, where possible, randomisation of sample order and blinding of experimenters handling samples. Data were analysed using R 4.0.2[42] and GraphPad Prism version 9 (GraphPad, San Diego).

## 3. Results

### 3.1 Characterization of undifferentiated knock-out MEFs

To assess the role of mitofusins in adipogenesis we utilised MEF lines deficient for Mfn1 (*Mfn1*^−/-^), Mfn2 (*Mfn2*^−/-^), or both (*Mfn1*^−/-^*2*^−/-^) (Figure 1A). We initially also characterised a line deficient for *Opa1* (*Opa1*^−/-^), the GTPase that mediates fusion of the inner mitochondrial membrane downstream of mitofusin engagement [43,44]. *Opa1*^−/-^ MEFs are expected to have a specific defect in mitochondrial fusion.

**Figure 1:**
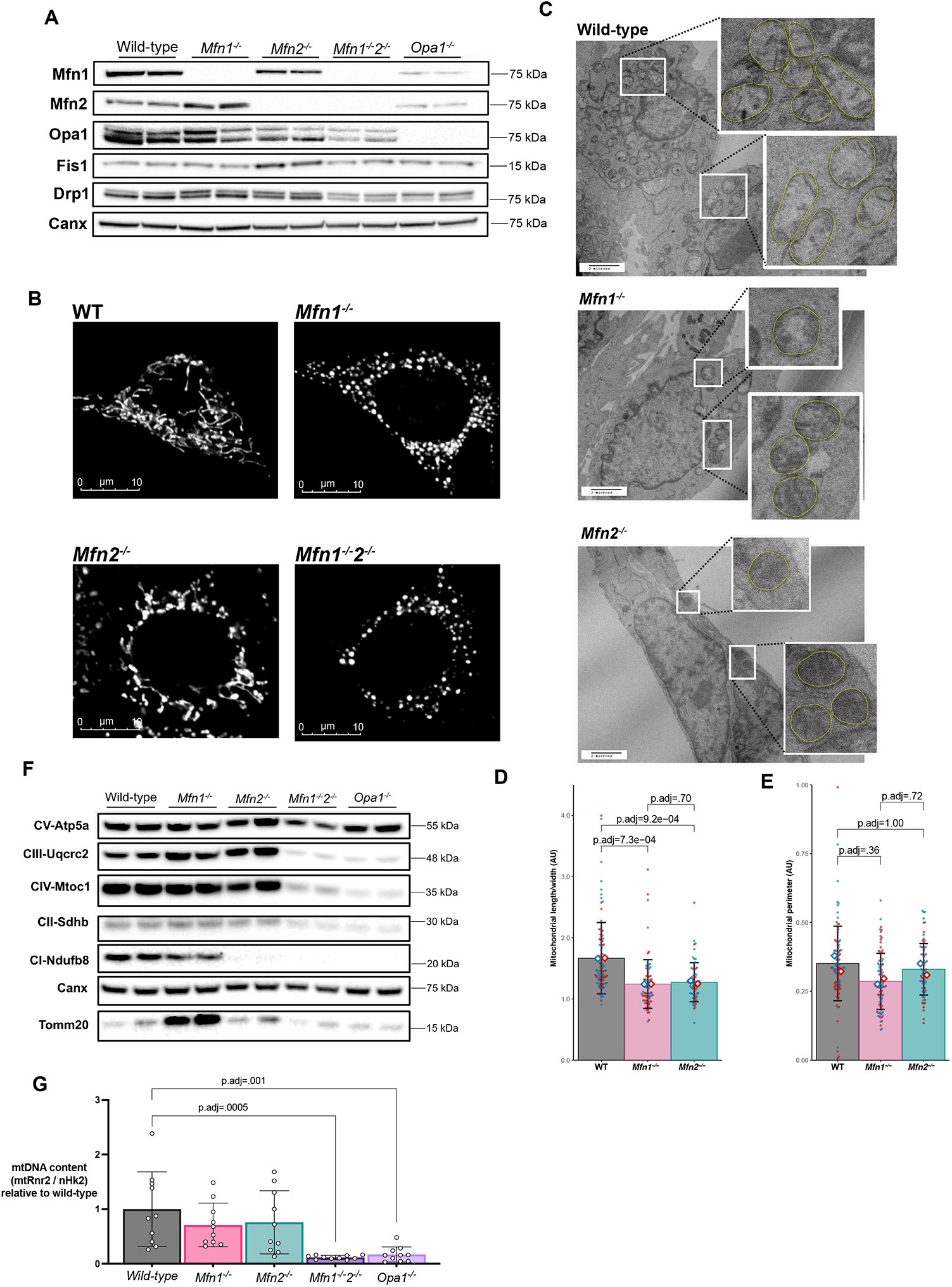
Loss of mitofusins induces mitochondrial network fragmentation in mouse embryonic fibroblasts (MEFs). MEFs deficient for the mitochondrial fusion proteins *Mfn1, Mfn2*, both *Mfn1* and *Mfn2* (*Mfn1*^−/-^*2*^−/-^), or *Opa1*^−/-^ were studied in their undifferentiated state. (A) Western blot confirming loss of protein from knock-out MEFs with calnexin (Canx) as loading control. (B) Mitotracker Orange imaging of MEFs on confocal microscopy demonstrated mitochondrial network fragmentation in knockout cell lines. (C) Transmission electron microscope (TEM) images of MEFs with zoomed-in images of mitochondria (highlighted in yellow). (D) Quantification of mitochondrial circularity (length/width) from TEM. Each red/blue dot represents one mitochondrion from two biological replicates. (E) Quantification of mitochondrial perimeter from TEM. Each red/blue dot represents one mitochondrion from two biological replicates. (F) Western blot of mitochondrial Oxphos complex subunits. (G) Mitochondrial DNA content expressed as mtRnr2 / nuclear Hk2 DNA from quantitative PCR. Each data point represents a separate biological replicate. All p-values represent pairwise comparisons between knock-outs and wild-type using t-tests, adjusted for multiple testing (p.adj). Data is representative of at least 3 independent replicates.

We first assessed expression of core proteins involved in mitochondrial fusion and fission in undifferentiated MEFs. Loss of *Mfn1* increased expression of Mfn2, whilst loss of *Mfn2* or *Opa1* reduced expression of Mfn1 (Figure 1A & Supplemental Fig. 1). Loss of each of the mitofusins was associated with no change in Opa1 expression whereas loss of both mitofusins led to a substantial reduction in Opa1 expression. Drp1, the main regulator of mitochondrial division [45], was expressed at similar levels in wild-type (WT) and *Mfn1*^−/-^, but increased in *Mfn2*^−/-^ MEFs and reduced in *Mfn1*^−/-^*2*^−/-^ and *Opa1*^*-/-*^ MEFs. Fis1, a protein involved in, but not required for, mitochondrial fission[46,47], was expressed at higher levels in *Mfn2*^−/-^ MEFs but at similar levels in the other MEF lines.

As previously reported[14], *Mfn1*^−/-^, *Mfn2*^−/-^ and *Mfn1*^−/-^*2*^−/-^ MEFs exhibited fragmented mitochondria when compared to WT controls using either confocal- (Figure 1B) or transmission electron-microscopy (TEM) (Figure 1C). Analysis of the TEM images revealed more circular mitochondria (reduced length/width) in *Mfn1*^−/-^ and *Mfn2*^−/-^ MEFs (Figure 1D) without any difference in the mean mitochondrial perimeter (Figure 1E). Mitochondrial network disruption was most striking in *Mfn1*^−/-^*2*^−/-^ MEFs, which also consistently multiplied very slowly and failed to show any signs of differentiation (data not shown). Growth of *Opa1*^−/-^ MEFs, which are also known to manifest severely disrupted mitochondrial morphology[48,49], was similarly delayed and they too failed to show any signs of adipocyte differentiation (data not shown), so these cells were not studied further in terms of adipogenesis. Loss of *Opa1* or both mitofusins has previously been found to prevent cells from responding to changes in metabolic demand[50] as well as manifesting low growth rates[43], which would limit the proliferation necessary for adipogenesis[51].

In the *Mfn1*^−/-^*2*^−/-^ and *Opa1*^−/-^ MEFs we observed substantially reduced expression of mitochondrial oxidative phosphorylation complexes II, III, and IV (Figure 1F & Supplemental Fig. 1). In contrast, Oxphos subunit expression was similar to WT cells in *Mfn1*^−/-^ MEFs, except for slightly increased complex III expression. TOMM20 expression was increased in *Mfn1*^−/-^ MEFs (Figure 1F & Supplemental Fig. 1), suggesting increased mitochondrial mass. In *Mfn2*^−/-^ cells expression of complexes II-V appeared normal whereas complex I expression was reduced (Figure 1F).

Mitochondrial DNA (mtDNA) is prone to depletion in cells with impaired mitochondrial fusion[52]. We found that double knock-out *Mfn1*^−/-^*2*^−/-^ and *Opa1*^−/-^ MEFs had reduced mtDNA levels whereas loss of *Mfn1* or *Mfn2* in isolation did not affect mtDNA content (Figure 1G).

### 3.2 Adipogenic differentiation of knock-out MEFs

We next assessed the ability of mitofusin KO MEFs to differentiate into adipocytes using a well-established adipogenic protocol. Surprisingly, we observed a specific increased lipid accumulation in *Mfn1*^−/-^ MEFs, but not in *Mfn2*^*-/-*^ MEFs, from day 4 of differentiation (Figure 2A), with increased neutral lipid content on both days 4 and 8 (Figure 2). A panel of mature adipocyte proteins, namely Plin1, Pparg, Fabp4, and Glut4 (Figure 2C & Supplemental Fig. 2) showed concomitant increases in expression. Loss of *Mfn1* was associated with a greater increase in Pparg1 than Pparg2 (Figure 2C & Supplemental Fig. 2). These expression changes corresponded to increased insulin-stimulated 2-deoxy-glucose uptake compared to WT MEFs (p=1.1×10^−12^, Figure 2D). These findings suggest *bona fide* enhanced adipogenic differentiation rather than simple lipid accumulation in *Mfn1*^*-/-*^ MEFs.

**Figure 2:**
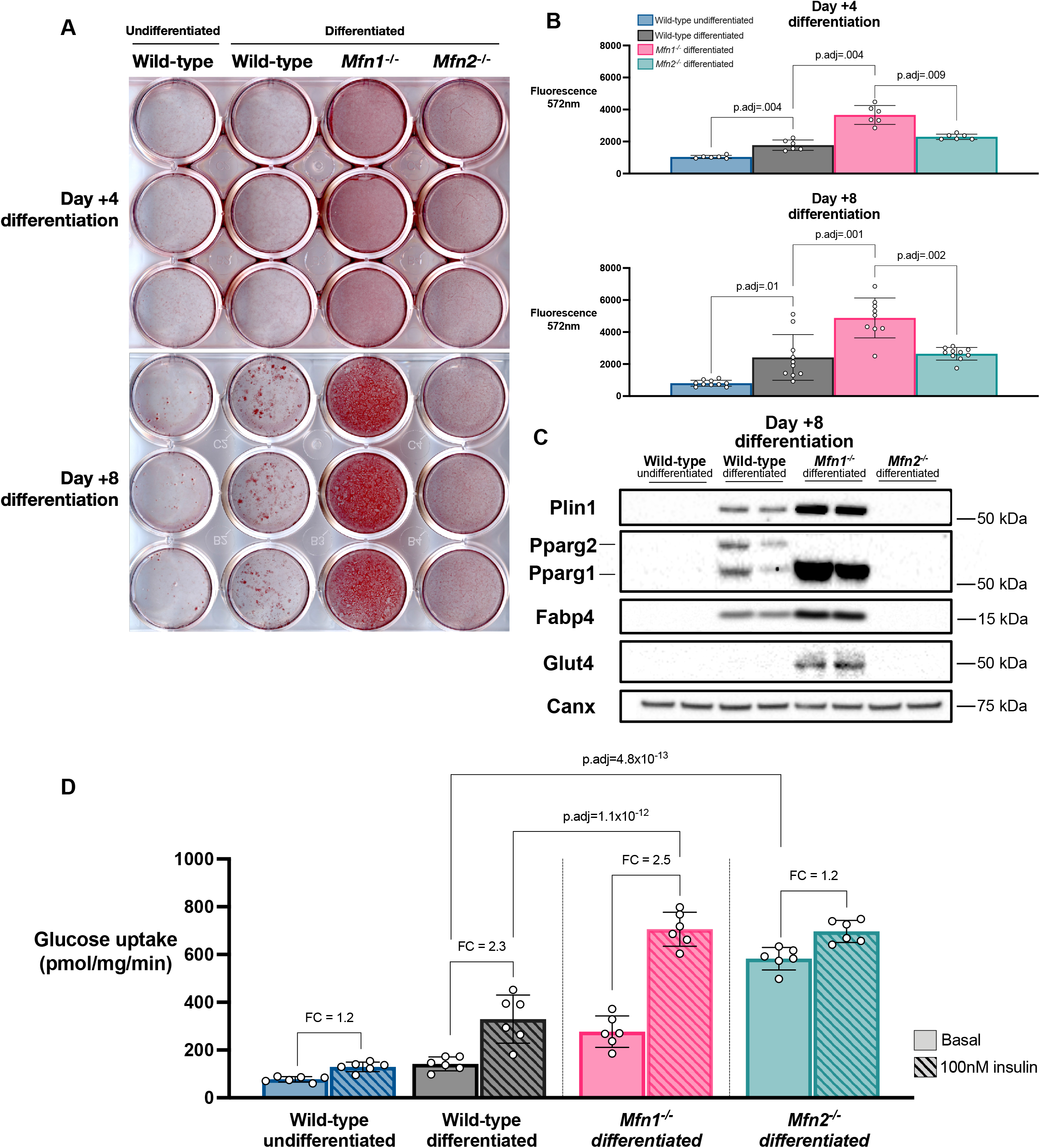
*Mfn1*^−/-^ MEFs demonstrate enhanced adipogenesis. MEFs underwent adipogenic differentiation using standard protocols and were assessed at day +4 and +8 of differentiation. (A) Oil Red O staining of differentiated MEFs compared to undifferentiated wild-type MEFs. (B) Fluorometric quantification of AdipoRed neutral lipid dye at day +4 & day +8. Each data point represents a separate biological experiment. (C) Western blot for markers of adipocyte differentiation at day +8. Calnexin (Canx) used as loading control. (D) Glucose uptake assay (2- deoxy-D-glucose) at day +8 under basal and 100nM insulin conditions. Each data point represents a separate biological experiment. FC = Fold increase from basal to insulin-stimulated glucose uptake. All p-values represent pairwise comparisons between knock-outs and wild-type using T-tests, adjusted for multiple testing (p.adj). Data is representative of at least 3 independent replicates.

Knockout of *Mfn2* had a contrasting effect on adipogenesis of MEFs. Lipid accumulation in differentiating *Mfn2*^−/-^ MEFs was similar to that of WT cells, however expression of Plin1, Pparg, and Fabp4 was lower than that in WT cells and *Glut4* expression was undetectable (Figure 2C & Supplemental Fig. 2). *Mfn2*^−/-^ MEFs also demonstrated minimal insulin-stimulated 2-deoxy-glucose uptake (1.2-fold change with 100nM insulin, Figure 2D), despite higher basal 2-deoxy-glucose uptake (WT 2.8 ± .6 vs. *Mfn2*-/- 11.7 ± 0.9 nmol/mg/min, p=4.8×10^−13^).

### 3.3 Differentiation of knock-out MEFs over-expressing Pparg

MEFs are not fully committed preadipocytes, and show a relatively low rate of adipocyte differentiation in response to hormonal stimuli alone [53]. We thus next assessed whether over- expression of *Pparg2*, the master transcriptional driver of adipogenesis, would modify the impact of mitofusins deficiency on adipogenesis. Pparg2 was retrovirally overexpressed in WT and different mitofusins KO MEFs, with overexpression higher in *Mfn2*^−/-^ MEFs than in WT and *Mfn1*^−/-^ MEFs (Supplemental Fig. 3A-B). Upon adipogenic differentiation, we observed results consistent with those in untransduced cells, namely enhanced lipid accumulation in *Mfn1*^−/-^ MEFs but similar lipid accumulation in the WT and *Mfn2*^−/-^ MEFs (Supplemental Fig. 3B & 3C). There was increased expression of *Fabp4* and *Glut4* in *Mfn1*^−/-^ MEFs with reduced expression of *Glut4* in *Mfn2*^−/-^ MEFs despite higher baseline expression of *Pparg2* (Supplemental Fig. 2 & 3D). Expression of Plin1 was higher in both *Mfn1*^−/-^ and *Mfn2*^−/-^ MEFs compared to differentiated WT MEFs, demonstrating that loss of *Mfn1*, but not *Mfn2*, was associated with enhanced differentiation.

### 3.4 Transcriptomic profiling of knock-out MEFs

To investigate how *Mfn1* deficiency enhances adipogenic differentiation we next performed bulk RNA sequencing of WT, *Mfn1*^−/-^, and *Mfn2*^−/-^ MEFs in undifferentiated (day -2) and differentiated (day +8) states. *Mfn1* and *Mfn2* null pre-adipocytes (day -2) both demonstrated significant differences to WT cells, and differences from each other (Figure 3A & 3B). Most strikingly, gene set enrichment analysis (GSEA) showed that the most highly enriched pathway in *Mfn1*^−/-^ MEFs was ‘hypoxia-related genes’ (Figure 3C), whereas this gene set was downregulated in *Mfn2*^−/-^ MEFs (Figure 3C). This gene set includes *Vegfa* as well as genes involved in the response to oxidative stress, such as *Selenbp1* and *Ankzf1*. Both knock-out lines showed enrichment of growth-related pathways (e.g. Estrogen Response Early), which was most evident in the *Mfn1*^−/-^ MEFs (Figure 3C). Genes implicated in the G2-M checkpoint gene set were upregulated in *Mfn2*^−/-^ MEFs but downregulated in *Mfn1*^−/-^ MEFs. *Mfn2*^−/-^ MEFs also showed upregulation of other pathways related to growth, including ‘Apoptosis’, ‘Mitotic spindle’, and ‘E2F targets’ (Figure 3C), consistent with the rapid growth of this cell line[54].

**Figure 3:**
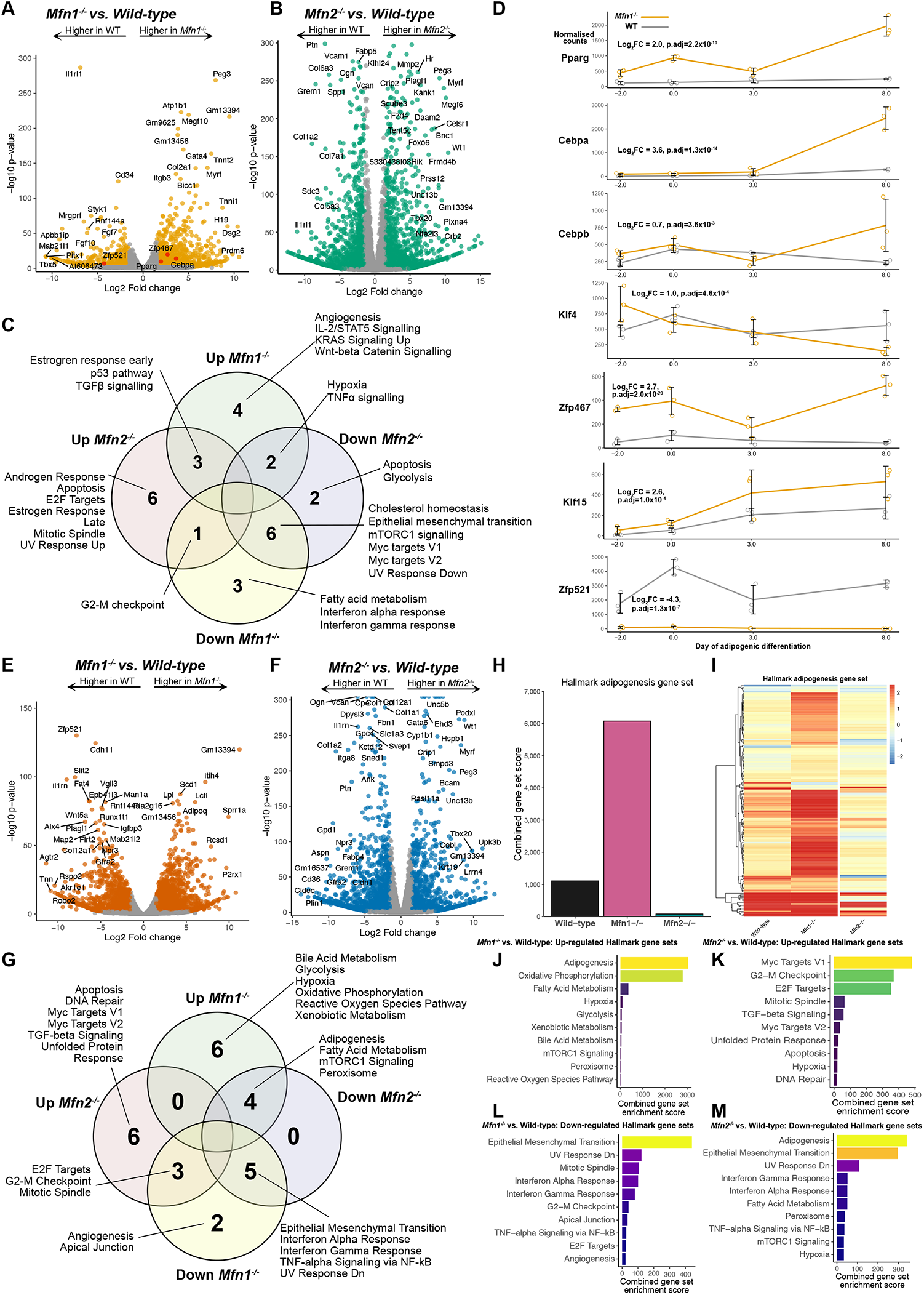
*Mfn1*^−/-^ MEFs demonstrate a pro-adipogenic transcriptional signature even in an undifferentiated state. Bulk RNA sequencing was performed on wild-type (WT), *Mfn1*^−/-^, and *Mfn2*^−/-^ at day -2, 0, +3, and +8 of differentiation. (A) Volcano plot demonstrating significant differential gene expression (DGE) in *Mfn1*^−/-^ MEFs vs. WT at day-2. Data from n=3 biological replicates. All genes in orange show significant DGE (adjusted p-value <0.001 and log_2_ fold change (log_2_FC) greater or less than 1.5. Adipogenic transcription factors are shown in red. (B) Volcano plot for *Mfn2*^−/-^ MEFs vs. WT at day -2. Genes in green show significant DGE. (C) Venn diagram illustrating the significantly upregulated and downregulated Hallmark gene sets on transcriptomic analysis of *Mfn1*^−/-^ and *Mfn2*^−/-^ MEFs versus wild-type MEFs at day -2. (D) Time course from RNAseq demonstrating the change in seven key adipogenic transcription factors in WT and *Mfn1*^−/-^ MEFs during differentiation. Data represents normalised counts per million. Log_2_FC and adjusted p-values (p.adj) are derived from calculations of DGE at day -2, as described above. (E) Volcano plot for *Mfn1*^−/-^ MEFs vs. WT at day+8. Data from n=3 biological replicates. All genes in red show significant DGE. (F) Volcano plot for *Mfn2*^−/-^ MEFs vs. WT at day +8. Genes in blue show significant DGE. (G) Significantly upregulated and downregulated Hallmark gene sets from *Mfn1*^−/-^ and *Mfn2*^−/-^ MEFs versus wild-type MEFs at day +8. (H) Hallmark adipogenesis gene set enrichment score from pathway analysis following DGE comparison of day -2 versus day +8 for each cell line. (I) Heatmap illustrating log2fold change for all the 200 genes included in the Hallmark adipogenesis gene set (data from (H)). Pathway analysis showing up-regulated (J-K) and down-regulated (L-M) pathways for *Mfn1*^−/-^ and *Mfn2*^−/-^ MEFs versus WT at day +8.

Although this analysis did not highlight the ‘adipogenesis pathway’, it did show that mRNA expression of some pro-adipogenic transcription factors, including *Pparg, Cebpa*, and *Zfp467*[55] was increased and that expression of *Zfp521* (coloured red in Figure 3A), an anti-adipogenic factor[56,57], was reduced in Mfn1^−/-^ cells compared to WT(Figure 3D). These differences were sustained throughout the differentiation time-course. Despite the higher basal glucose uptake in *Mfn2*^−/-^ MEFs compared to WT (Figure 2D), there was no significant difference in *Slc2a1* (gene encoding Glut1): 0.2 log2FC, q-value = .03. Unlike models of *Mfn2* deletion *in vivo*[11], there was no evidence of activation of the integrated stress response in *Mfn2*^*-/-*^ MEFs in the undifferentiated state (Figure 3C).

RNA sequencing of mature (day +8) adipocytes (Figure 3E-F) demonstrated expression profiles consistent with enhanced differentiation in *Mfn1*^−/-^ MEFs and reduced differentiation in *Mfn2*^−/-^ MEFs. For example, relative to WT, *Mfn1*^−/-^ MEFs showed increased expression of *Plin1, Pparg, Slc2a4* (encoding Glut4), *Fabp4*, and *Cd36*, whilst each of these were reduced in *Mfn2*^−/-^ MEFs. This was also observed on gene set enrichment analysis using Hallmark gene sets[41]: when comparing day -2 pre-adipocytes and day +8 mature adipocytes within each cell line, *Mfn1*^*-/-*^ had markedly greater enrichment of the adipogenesis gene set (Figure 3H-I). Similarly, pathway analysis comparing day +8 WT and KO cell lines revealed upregulation of the adipogenesis gene set for *Mfn1*^−/-^ MEFs and downregulation of the same gene set in *Mfn2*^−/-^ MEFs (Figure G-M). *Mfn1*^−/-^ MEFs also manifested upregulation of oxidative phosphorylation, hypoxia, and reactive oxygen species pathway gene sets (Figure 3J). Thus, the transcriptomic data supports a pro-adipogenic phenotype in *Mfn1*^−/-^, but not *Mfn2*^−/-^, MEFs.

### 3.5 siRNA knockdown and differentiation of 3T3-L1 preadipocytes

Given the surprisingly divergent effects of Mfn1 and Mfn2 deficiencies on MEF adipogenesis, we next sought to corroborate findings in an independent cellular model. We thus evaluated the impact of siRNA mediated *Mfn1* and *Mfn2* knockdown in 3T3-L1 pre-adipocytes. The 3T3-L1 line is a MEF subclone first established in the 1970s based on its high propensity for adipogenic differentiation, and has since been widely used as a model of *in vitro* adipogenesis[58,59].

Treating 3T3-L1s with siRNAs targeting *Mfn1* or *Mfn2* effectively reduced target protein expression by more than 95% (Figure 4A and Supplementary Fig. 4A-B). Like the KO cell lines, knock-down of Mfn2 reduced Mfn1 expression, whereas knock-down of Mfn1 modestly increased Mfn2 expression (Figure 4A & Supplementary Fig. 4).

**Figure 4:**
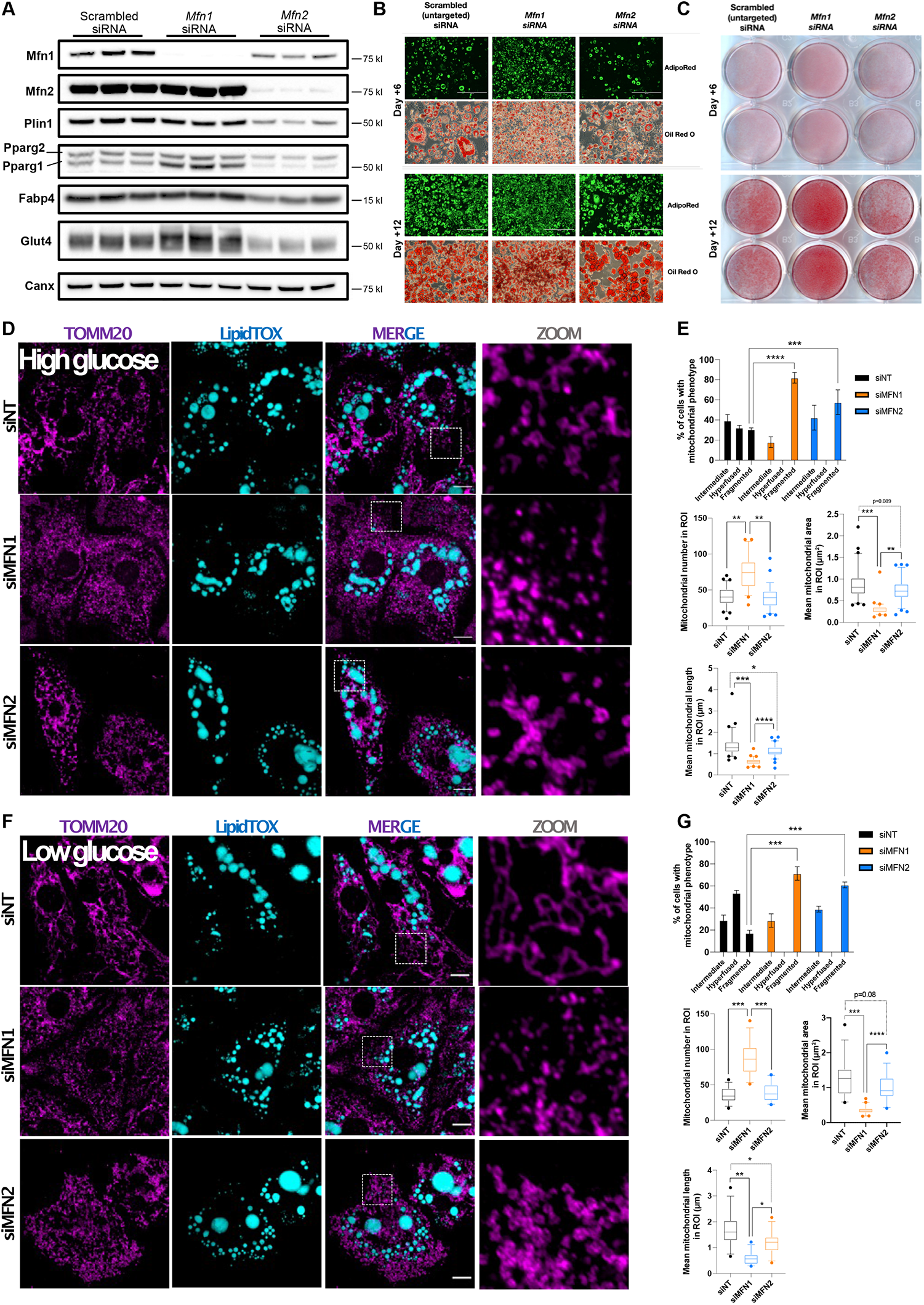
siRNA knockdown of *Mfn1* in 3T3-L1s enhances adipogenesis. 3T3-L1s were treated with scrambled, *Mfn1*-targeted, or *Mfn2*-targeted siRNA on alternate days from day -2 to day +12 of differentiation and assessed on days +6 and +12. (A) Western blot from 3T3-L1 at day +12 showing efficacy of *Mfn1*/*Mfn2* knock-down and expression of selected adipogenic genes. Data represents three biological replicates. (B) Light microscope images of AdipoRed fluorescence (green=lipid) and Oil Red O stained 3T3-L1s at day +6 and +12 of differentiation. (C) Oil Red O staining of 3T3-L1s at day +6 and +12 of differentiation. (D) Effect of siRNA knock-down on mitochondrial morphology assessed by confocal microscopy in 3T3-L1s cells at day +12 of differentiation under high-glucose conditions. Mitochondria were labelled using an anti-TOMM20 antibody (purple) and neutral lipids were stained with LipidTox (blue). Scale bars: 10 μm. (E) Quantification of mitochondrial morphology, number, area, and length in 225 μm^2^ region of interests (ROI) from (D). (F) Effect of siRNA knock-down on mitochondrial morphology assessed by confocal microscopy in 3T3-L1s cells at day +12 of differentiation under under low glucose conditions. Mitochondria were labelled using an anti-TOMM20 antibody (purple) and neutral lipids were stained with LipidTox (blue). Scale bars: 10 μm. (G) Quantification of mitochondrial morphology, number, area, and length in 225 μm^2^ ROIs from (F). All p-values represent pairwise comparisons between knock-outs and wild-type using t-tests, adjusted for multiple testing (p.adj). Data is representative of at least 3 independent replicates.

In keeping with the MEF data, 3T3-L1s treated with *Mfn1*-targeted siRNA demonstrated increased lipid accumulation (Figure 4B-C). Whilst there was no difference in lipid accumulation between scrambled and *Mfn2* knockdown cells, there was a trend towards increased lipid droplet size (Figure 4F). Expression of Pparg1 and Glut4 protein was increased in *Mfn1* knockdown cells (Figure 4A & Supplementary Fig. 4), whereas expression of Plin1 and Fabp4 were similar to that of the control cells. *Mfn2* knock-down reduced lipid accumulation and expression of Plin1, Fabp4, Pparg1, Pparg2, and Glut4 proteins (Figure 4 & Supplementary Fig. 4).

Mitochondrial morphology analysis revealed that loss of Mfn1 and Mfn2 both led to mitochondrial fragmentation, with Mfn1 silencing inducing a more drastic changes of the mitochondrial network in mature (day +12) differentiated adipocytes (Figure 4D-E), even though, as previously reported[25], the mitochondrial network of wild-type 3T3-L1s manifests increased fragmentation in late differentiation (Figure 4D-E). Nutritional deprivation is known to induce mitochondrial fusion in several different cell lines[60]. In keeping with this, incubating the 3T3-L1 adipocytes in low glucose media or without serum increased fusion of the mitochondrial network in siNT treated control cells characterized by mitochondrial elongation (Supplementary Fig. 5), accentuating the impact of Mfn1 and Mfn2 deficiencies on the mitochondrial network (Figure 4F-G). These findings in 3T3-L1 adipocytes support the observation that loss of two mitofusins have specific effects of adipogenic potential despite both causing mitochondrial network fragmentation.

## 4. Discussion

Prompted by observations of a human monogenic disease, which highlighted the importance of the mitochondrial network in adipose tissue, we sought to investigate the roles of two mitochondrial fusion proteins in adipogenesis. Although human genetics has established that a specific genetic mutation of *MFN2* has a profound impact on white adipose tissue development and function, our analysis of adipogenesis in cultured cells suggested that deletion or knockdown of *Mfn1* enhances adipogenesis whereas *Mfn2* deletion or knockdown tended to impair differentiation.

The absence of Mfn2 in MEFs subtly reduced adipogenesis, as manifest by reduced expression of Plin1, Pparg, and Glut4, with no change in neutral lipid accumulation. *Mfn2*^−/-^ adipocytes had increased basal glucose uptake but a significantly impaired response to insulin stimulation. This may relate to GLUT1 expression in the basal state and very low or absent GLUT4 expression. Data from tissue-specific knock-out mice suggests that loss of *Mfn2* can perturb glucose homeostasis. Specifically, deletion of *Mfn2* in all adipocytes[20] or in brown adipocytes alone[21] caused reduced expression of multiple oxidative phosphorylation subunits and impaired cold tolerance yet, paradoxically, both mouse lines were protected from systemic insulin resistance. Mice in which Mfn2 was inducibly deleted in all adipocytes in adulthood showed increased obesity and elevated blood glucose[22].

In both *Mfn1*^*-/-*^ knock-out MEFs and *Mfn1* knock-down 3T3-L1s there was a significant increase in lipid accumulation. In our view, this reflects enhanced adipogenesis, as it was associated with elevated expression of several stereotypical adipogenic markers of (e.g. Plin1, FABP4 and *Glut4*). *Mfn1*^*-/-*^ MEFs had increased Pparg expression even in the undifferentiated state, and RNA sequencing showed that expression of some other pro-adipogenic transcription factors (*Cebpa, Cebpb, Klf4, Klf15*, and *Zfp467*) was increased, suggesting that Mfn1 deficiency may prime cells transcriptionally to favour adipogenesis. Indeed, the changes in gene expression observed during differentiation showed a stronger signature for adipogenic differentiation in Mfn1^−/-^ than wild-type MEFs.

The exact mechanism behind this is currently unclear. One potential explanation could involve the generation of oxidised lipids secondary to ROS generation from altered mitochondrial function. This is a plausible hypothesis for these observations as increased ROS early in adipogenesis has been reported to accelerate differentiation[61]. In addition, humans with loss of function of selenoproteins have chronically elevated oxidative stress and have increased adiposity yet are more insulin sensitive[62]. Studies of ROS stimulation and blockade in various models of mitochondrial dysfunction may help to delineate a causal role for ROS in the enhanced adipogenesis observed in *Mfn1*-deficient cells. However, such experiments require careful titration of relatively toxic agents that may inhibit adipogenesis through non-specific cellular damage. Multiple tissue-specific knock-outs for both *Mfn1*[63] and *Mfn2*[12,64] have identified changes in ROS. Liver-specific knock-out of *Mfn2* was associated with increased hydrogen peroxide and reduced expression of subunits from Complex IV[12]. In addition, *Mfn2*^−/-^ MEFs have reduced membrane potential compared to wild-type[65], which, along with lower Complex I expression, may influence their degree for ROS generation through reverse electron transport[66,67].

An important limitation of this work is a lack of *in vivo* data to support this hypothesised role for *Mfn1* in adipogenesis. Adipose-specific *Mfn1* knock-out mice, although briefly said in one study not to have a gross adipose phenotype, have not been characterised in any detail[20]. An important limitation of such a model in any case relates to the fact that deletion of ‘floxed’ *Mfn1* was under control of the adiponectin-promoter[20]. Adiponectin is only expressed in mature adipocytes[68] so such an experiment does not test the impact of Mfn1 deficiency early in adipocyte development. The data reported herein was also exclusively conducted in murine cells. Whilst we found broadly concordant observations in two different cell lines we have not studied human adipocytes. There are currently no known human disorders linked to *MFN1* mutations but based on our findings, we would hypothesise that individuals carrying such mutations may be more insulin sensitive and therefore may not present with a clinical disease phenotype. However, germline *Mfn1* mutations would also perturb Mfn1 function in other tissues which may have a greater phenotypic impact than that related to altered adipose expression. On the other hand human population frequencies of sequence variants in *MFN1* and *MFN2* suggest that loss-of-function mutations in *MFN1* may not be selected against, unlike loss-of-function variants in *MFN2* [69].

### 4.1 Conclusions

At endogenous levels, MFN1 appears to exercise tonic physiological restraint on differentiation to adipocytes in culture, whereas the closely related MFN2 is necessary for expression of the full programme of adipogenesis.

## Abbreviations^1^

^1^

2DOG: 2-deoxy-glucose
BDMA: Benzyldimethylamine
BSA: bovine serum albumin
DGE: Differential gene expression
DMEM: Dulbecco’s Modified Eagle’s Medium
DIW: deionised water
FBS: foetal bovine serum
FDR: false-discovery rate
IBMX: 3-Isobutyl-1-methylxanthine
MEF: mouse embryonic fibroblast
Mfn1/2: Mitofusin 1/2
mtDNA: mitochondrial DNA
ORO: oil red O
Opa1: Optic atrophy 1
PBS: phosphate buffered saline
Pparg: Peroxisome proliferator-activated receptor gamma
ROS: reactive oxygen species
RT-qPCR: real-time quantitative polymerase chain reaction
TEM: Transition electron microscope
WT: wild-type.

## 5. Acknowledgements

We are grateful for the support of the Cambridge Advanced Imaging Centre (University of Cambridge, UK) for performing, processing, and imaging of transmission electron microscope studies. We thank M. Mimmack for his technical assistance with cell culture and assays.

## Supplementary Data

**Supplementary Figure 1**: **Quantification of protein expression from immunoblots in Figure 1**. Quantification of protein expression (relative to wild-type) for all proteins shown in figures 1A and 1F. Each data point represents a separate biological replicate. Stars indicate p-values following pairwise comparisons between groups: * p<0.5, ** p<.01, *** p<.001, **** p<.0001. ns, not significant.

**Supplementary Figure 2**: **Quantification of protein expression from immunoblots for differentiation of knock-out MEFs**. (A-E) Quantification of protein expression (relative to wild-type) for all proteins shown in figures 2C from knock-out MEFs. (F-J) Quantification of protein expression from knock-out MEFs over-expressing Pparg2, from Supplementary Figure 3D. Each data point represents a separate biological replicate. Stars indicate p-values following pairwise comparisons between groups: * p<0.5, ** p<.01, *** p<.001, **** p<.0001. ns, not significant.

**Supplementary Figure 3**: **Increased adipogenesis in *Mfn1***^**-/-**^ **MEFs even in setting of *Pparg2* over-expression**. *Pparg2* was over-expressed in MEFs to enhance their adipogenic differentiation capacity and cells were assessed on day +4 and +8 of protocols. (A) Western blot illustrating the relative Pparg2 over-expression in cell lines used compared to non-transduced wild-type cells. (B) Quantification of Pparg2 expression from the Western blot in panel (A). (C) Oil Red O staining differentiated MEFs compared to undifferentiated wild-type MEFs. (D) Fluorometric quantification of AdipoRed neutral lipid dye at day +4 & day +8. Each data point represents a separate biological experiment. Data is expressed relative to undifferentiated wild-type for each biological replicate. (E) Western blot for markers of adipocyte differentiation at day +8 from MEFs over-expressing *Pparg2*. All p-values represent pairwise comparisons between knock-outs and wild-type using t-tests, adjusted for multiple testing (p.adj). Data is representative of at least 3 independent replicates.

**Supplementary Figure 4**: **Quantification of protein expression from immunoblots in Figure 4A**. Quantification of protein expression (relative to scrambled control) for all proteins shown in figures 4A. Each data point represents a separate biological replicate. Stars indicate p-values following pairwise comparisons between groups: * p<0.5, ** p<.01, *** p<.001, **** p<.0001. ns, not significant.

**Supplementary Figure 5**: **Low glucose treated induces mitochondrial network fusion in differentiated mature 3T3-L1 adipocytes**. (A) Representative confocal images of 3T3-L1s cells at day +12 of differentiation under high-glucose (HG) and low glucose (LG) conditions. Mitochondria were labelled using an anti-TOMM20 antibody (purple) and neutral lipids were stained with LipidTox (blue). Scale bars: 10 μm. (B) Quantification of mitochondrial morphology, number, area, and length in 225 μm^2^ region of interests from (A). Stars indicate p-values following pairwise comparisons between groups: * p<0.5, *** p<.001.

**Supplementary Table 1**: **Reagents and cell lines**. List of all reagents and equipment used in this study.

**Supplementary Table 2**: **Antibodies used**. List of all primary and secondary antibodies used in this study.

**Supplementary Data 1. RNA sequencing counts tables**. Normalised counts per million per gene from RNA sequencing of MEFs. dm2, day -2 of differentiation; dm0, day zero of differentiation; dp3, day +3 of differentiation; dp8, day +8 of adipogenic differentiation; m1, Mfn1-/-; m2, Mfn2-/-; WT, wild-type MEFs.

## References

[1] Gorman, G.S., Chinnery, P.F., DiMauro, S., Hirano, M., Koga, Y., McFarland, R., et al., 2016. Mitochondrial diseases. Nature Reviews Disease Primers 2: 16080.

[2] Wallace, D.C., 2018. Mitochondrial genetic medicine. Nature Genetics 50(12): 1642–9.

[3] Mann, J.P., Savage, D.B., 2019. What lipodystrophies teach us about the metabolic syndrome. The Journal of Clinical Investigation 130: 4009–21.

[4] Rutkowski, J.M., Stern, J.H., Scherer, P.E., 2015. The cell biology of fat expansion. The Journal of Cell Biology 208(5): 501–12.

[5] Rocha, N., Bulger, D.A., Frontini, A., Titheradge, H., Gribsholt, S.B., Knox, R., et al., 2017. Human biallelic MFN2 mutations induce mitochondrial dysfunction, upper body adipose hyperplasia, and suppression of leptin expression. ELife 6: e23813.

[6] Sawyer, S.L., Ng, A.C.H., Micheil Innes, A., Wagner, J.D., Dyment, D.A., Tetreault, M., et al., 2015. Homozygous mutations in MFN2 cause multiple symmetric lipomatosis associated with neuropathy. Human Molecular Genetics 24(18): 5109–14.

[7] Capel, E., Vatier, C., Cervera, P., Stojkovic, T., Disse, E., Cottereau, A.S., et al., 2018. MFN2-associated lipomatosis: Clinical spectrum and impact on adipose tissue. Journal of Clinical Lipidology, doi: 10.1016/j.jacl.2018.07.009.

[8] Pipis, M., Feely, S.M.E., Polke, J.M., Skorupinska, M., Perez, L., Shy, R.R., et al., 2021. Natural history of Charcot-Marie-Tooth disease type 2A: a large international multicentre study. Brain: A Journal of Neurology, doi: 10.1093/brain/awaa323.

[9] Mishra, P., Chan, D.C., 2016. Metabolic regulation of mitochondrial dynamics. The Journal of Cell Biology 212(4): 379–87.

[10] Tilokani, L., Nagashima, S., Paupe, V., Prudent, J., 2018. Mitochondrial dynamics: overview of molecular mechanisms. Essays in Biochemistry 62(3): 341–60.

[11] Liesa, M., Shirihai, O.S., 2013. Mitochondrial dynamics in the regulation of nutrient utilization and energy expenditure. Cell Metabolism 17(4): 491–506.

[12] Sebastian, D., Hernandez-Alvarez, M.I., Segales, J., Sorianello, E., Munoz, J.P., Sala, D., et al., 2012. Mitofusin 2 (Mfn2) links mitochondrial and endoplasmic reticulum function with insulin signaling and is essential for normal glucose homeostasis. Proceedings of the National Academy of Sciences 109(14): 5523–8.

[13] Willems, P.H.G.M., Rossignol, R., Dieteren, C.E.J., Murphy, M.P., Koopman, W.J.H., 2015. Redox Homeostasis and Mitochondrial Dynamics. Cell Metabolism 22(2): 207–18.

[14] Chen, H., Detmer, S.A., Ewald, A.J., Griffin, E.E., Fraser, S.E., Chan, D.C., 2003. Mitofusins Mfn1 and Mfn2 coordinately regulate mitochondrial fusion and are essential for embryonic development. The Journal of Cell Biology 160(2): 189–200.

[15] Santel, A., Frank, S., Gaume, B., Herrler, M., Youle, R.J., Fuller, M.T., 2003. Mitofusin-1 protein is a generally expressed mediator of mitochondrial fusion in mammalian cells. Journal of Cell Science 116(Pt 13): 2763–74.

[16] Rojo, M., Legros, F., Chateau, D., Lombes, A., 2002. Membrane topology and mitochondrial targeting of mitofusins, ubiquitous mammalian homologs of the transmembrane GTPase Fzo. Journal of Cell Science 115(Pt 8): 1663–74.

[17] de Brito, O.M., Scorrano, L., 2008. Mitofusin 2 tethers endoplasmic reticulum to mitochondria. Nature 456(7222): 605–10.

[18] Benador, I.Y., Veliova, M., Liesa, M., Shirihai, O.S., 2019. Mitochondria Bound to Lipid Droplets: Where Mitochondrial Dynamics Regulate Lipid Storage and Utilization. Cell Metabolism 29(4): 827– 35.

[19] Chen, Y., Dorn, G.W., 2013. PINK1-phosphorylated mitofusin 2 is a Parkin receptor for culling damaged mitochondria. Science 340(6131): 471–5.

[20] Boutant, M., Kulkarni, S.S., Joffraud, M., Ratajczak, J., Valera-Alberni, M., Combe, R., et al., 2017. Mfn2 is critical for brown adipose tissue thermogenic function. The EMBO Journal 41: e201694914.

[21] Mahdaviani, K., Benador, I.Y., Su, S., Gharakhanian, R.A., Stiles, L., Trudeau, K.M., et al., 2017. Mfn2 deletion in brown adipose tissue protects from insulin resistance and impairs thermogenesis. EMBO Reports 18(7): e201643827.

[22] Mancini, G., Pirruccio, K., Yang, X., Blüher, M., Rodeheffer, M., Horvath, T.L., 2019. Mitofusin 2 in Mature Adipocytes Controls Adiposity and Body Weight. Cell Reports 26(11): 2849-2858.e4.

[23] Wilson-Fritch, L., Nicoloro, S., Chouinard, M., Lazar, M.A., Chui, P.C., Leszyk, J., et al., 2004. Mitochondrial remodeling in adipose tissue associated with obesity and treatment with rosiglitazone. The Journal of Clinical Investigation 114(9): 1281–9.

[24] Lu, R.-H., Ji, H., Chang, Z.-G., Su, S.-S., Yang, G.-S., 2010. Mitochondrial development and the influence of its dysfunction during rat adipocyte differentiation. Molecular Biology Reports 37(5): 2173–82.

[25] Ducluzeau, P.H., Priou, M., Weitheimer, M., Flamment, M., Duluc, L., Iacobazi, F., et al., 2011. Dynamic regulation of mitochondrial network and oxidative functions during 3T3-L1 fat cell differentiation. Journal of Physiology and Biochemistry 67(3): 285–96.

[26] Tontonoz, P., Hu, E., Spiegelman, B.M., 1994. Stimulation of adipogenesis in fibroblasts by PPARγ2, a lipid-activated transcription factor. Cell 79(7): 1147–56.

[27] Tang, Q.-Q., Otto, T.C., Lane, M.D., 2003. CCAAT/enhancer-binding protein beta is required for mitotic clonal expansion during adipogenesis. Proceedings of the National Academy of Sciences of the United States of America 100(3): 850–5.

[28] Mackall, J.C., Student, A.K., Polakis, E., Lane, D.M., 1976. Induction of lipogenesis during differentiation in a “preadipocyte” cell line. The Journal of Biological Chemistry 251(20): 6462–4.

[29] Nagashima, S., Tábara, L.-C., Tilokani, L., Paupe, V., Anand, H., Pogson, J.H., et al., 2020. Golgi-derived PI(4)P-containing vesicles drive late steps of mitochondrial division. Science 367(6484): 1366–71.

[30] Zhang, Y., Lanjuin, A., Chowdhury, S.R., Mistry, M., Silva-García, C.G., Weir, H.J., et al., 2019. Neuronal TORC1 modulates longevity via AMPK and cell nonautonomous regulation of mitochondrial dynamics in C. elegans. ELife 8, doi: 10.7554/eLife.49158.

[31] Lord, S.J., Velle, K.B., Mullins, R.D., Fritz-Laylin, L.K., 2020. SuperPlots: Communicating reproducibility and variability in cell biology. The Journal of Cell Biology 219(6), doi: 10.1083/jcb.202001064.

[32] Fazakerley, D.J., Chaudhuri, R., Yang, P., Maghzal, G.J., Thomas, K.C., Krycer, J.R., et al., 2018. Mitochondrial CoQ deficiency is a common driver of mitochondrial oxidants and insulin resistance. ELife 7: e32111.

[33] Martin, M., 2011. Cutadapt removes adapter sequences from high-throughput sequencing reads. EMBnet.Journal 17(1): 10–2.

[34] Dobin, A., Davis, C.A., Schlesinger, F., Drenkow, J., Zaleski, C., Jha, S., et al., 2013. STAR: ultrafast universal RNA-seq aligner. Bioinformatics 29(1): 15–21.

[35] Danecek, P., Bonfield, J.K., Liddle, J., Marshall, J., Ohan, V., Pollard, M.O., et al., 2021. Twelve years of SAMtools and BCFtools. GigaScience 10(2), doi: 10.1093/gigascience/giab008.

[36] Liao, Y., Smyth, G.K., Shi, W., 2014. featureCounts: an efficient general purpose program for assigning sequence reads to genomic features. Bioinformatics 30(7): 923–30.

[37] Love, M.I., Huber, W., Anders, S., 2014. Moderated estimation of fold change and dispersion for RNA-seq data with DESeq2. Genome Biology 15(12): 550.

[38] Xie, Z., Bailey, A., Kuleshov, M.V., Clarke, D.J.B., Evangelista, J.E., Jenkins, S.L., et al., 2021. Gene Set Knowledge Discovery with Enrichr. Current Protocols 1(3): e90.

[39] Chen, E.Y., Tan, C.M., Kou, Y., Duan, Q., Wang, Z., Meirelles, G.V., et al., 2013. Enrichr: interactive and collaborative HTML5 gene list enrichment analysis tool. BMC Bioinformatics 14: 128.

[40] Kuleshov, M.V., Jones, M.R., Rouillard, A.D., Fernandez, N.F., Duan, Q., Wang, Z., et al., 2016. Enrichr: a comprehensive gene set enrichment analysis web server 2016 update. Nucleic Acids Research 44(W1): W90–7.

[41] Liberzon, A., Birger, C., Thorvaldsdóttir, H., Ghandi, M., Mesirov, J.P., Tamayo, P., 2015. The Molecular Signatures Database (MSigDB) hallmark gene set collection. Cell Systems 1(6): 417–25.

[42] R Core Team., 2019. A language and environment for statistical computing. Vienna, Austria: R Foundation for Statistical Computing.

[43] Chen, H., Chomyn, A., Chan, D.C., 2005. Disruption of fusion results in mitochondrial heterogeneity and dysfunction. The Journal of Biological Chemistry 280(28): 26185–92.

[44] Davies, V.J., Hollins, A.J., Piechota, M.J., Yip, W., Davies, J.R., White, K.E., et al., 2007. Opa1 deficiency in a mouse model of autosomal dominant optic atrophy impairs mitochondrial morphology, optic nerve structure and visual function. Human Molecular Genetics 16(11): 1307–18.

[45] Smirnova, E., Griparic, L., Shurland, D.L., van der Bliek, A.M., 2001. Dynamin-related protein Drp1 is required for mitochondrial division in mammalian cells. Molecular Biology of the Cell 12(8): 2245–56.

[46] Losón, O.C., Song, Z., Chen, H., Chan, D.C., 2013. Fis1, Mff, MiD49, and MiD51 mediate Drp1 recruitment in mitochondrial fission. Molecular Biology of the Cell 24(5): 659–67.

[47] Otera, H., Wang, C., Cleland, M.M., Setoguchi, K., Yokota, S., Youle, R.J., et al., 2010. Mff is an essential factor for mitochondrial recruitment of Drp1 during mitochondrial fission in mammalian cells. The Journal of Cell Biology 191(6): 1141–58.

[48] Song, Z., Ghochani, M., McCaffery, J.M., Frey, T.G., Chan, D.C., 2009. Mitofusins and OPA1 mediate sequential steps in mitochondrial membrane fusion. Molecular Biology of the Cell 20(15): 3525–32.

[49] Song, Z., Chen, H., Fiket, M., Alexander, C., Chan, D.C., 2007. OPA1 processing controls mitochondrial fusion and is regulated by mRNA splicing, membrane potential, and Yme1L. The Journal of Cell Biology 178(5): 749–55.

[50] Patten, D.A., Wong, J., Khacho, M., Soubannier, V., Mailloux, R.J., Pilon-Larose, K., et al., 2014. OPA1-dependent cristae modulation is essential for cellular adaptation to metabolic demand. The EMBO Journal 33(22): 2676–91.

[51] Tang, Q.-Q., Otto, T.C., Lane, M.D., 2003. Mitotic clonal expansion: a synchronous process required for adipogenesis. Proceedings of the National Academy of Sciences of the United States of America 100(1): 44–9.

[52] Chen, H., Vermulst, M., Wang, Y.E., Chomyn, A., Prolla, T.A., McCaffery, J.M., et al., 2010. Mitochondrial fusion is required for mtdna stability in skeletal muscle and tolerance of mtDNA mutations. Cell 141(2): 280–9.

[53] Rosen, E.D., Sarraf, P., Troy, A.E., Bradwin, G., Moore, K., Milstone, D.S., et al., 1999. PPAR gamma is required for the differentiation of adipose tissue in vivo and in vitro. Molecular Cell 4(4): 611–7.

[54] Chen, K.H., Dasgupta, A., Ding, J., Indig, F.E., Ghosh, P., L. Longo D., 2014. Role of mitofusin 2 (Mfn2) in controlling cellular proliferation. FASEB Journal: Official Publication of the Federation of American Societies for Experimental Biology 28(1): 382–94.

[55] Quach, J.M., Walker, E.C., Allan, E., Solano, M., Yokoyama, A., Kato, S., et al., 2011. Zinc finger protein 467 is a novel regulator of osteoblast and adipocyte commitment. The Journal of Biological Chemistry 286(6): 4186–98.

[56] Addison, W.N., Fu, M.M., Yang, H.X., Lin, Z., 2014. Direct transcriptional repression of Zfp423 by Zfp521 mediates a bone morphogenic protein-dependent osteoblast versus adipocyte lineage commitment switch. And Cellular Biology.

[57] Kang, S., Akerblad, P., Kiviranta, R., Gupta, R.K., Kajimura, S., Griffin, M.J., et al., 2012. Regulation of early adipose commitment by Zfp521. PLoS Biology 10(11): e1001433.

[58] Todaro, G.J., Green, H., 1963. Quantitative studies of the growth of mouse embryo cells in culture and their development into established lines. The Journal of Cell Biology 17: 299–313.

[59] Rubin, C.S., Hirsch, A., Fung, C., Rosen, O.M., 1978. Development of hormone receptors and hormonal responsiveness in vitro. Insulin receptors and insulin sensitivity in the preadipocyte and adipocyte forms of 3T3-L1 cells. The Journal of Biological Chemistry 253(20): 7570–8.

[60] Rambold, A.S., Kostelecky, B., Elia, N., Lippincott-Schwartz, J., 2011. Tubular network formation protects mitochondria from autophagosomal degradation during nutrient starvation. Proceedings of the National Academy of Sciences 108(25): 10190–5.

[61] Tormos, K.V., Anso, E., Hamanaka, R.B., Eisenbart, J., Joseph, J., Kalyanaraman, B., et al., 2011. Mitochondrial complex III ROS regulate adipocyte differentiation. Cell Metabolism 14(4): 537–44.

[62] Schoenmakers, E., Gurnell, M., Chatterjee, K., Schoenmakers, E., Agostini, M., Mitchell, C., et al., 2010. Mutations in the selenocysteine insertion sequence – binding protein 2 gene lead to a multisystem selenoprotein deficiency disorder in humans. The Journal of Clinical Investigation 120(12): 4220–35.

[63] Ramírez, S., Gómez-Valadés, A.G., Schneeberger, M., Varela, L., Haddad-Tóvolli, R., Altirriba, J., et al., 2017. Mitochondrial Dynamics Mediated by Mitofusin 1 Is Required for POMC Neuron Glucose-Sensing and Insulin Release Control. Cell Metabolism 25(6): 1390-1399.e6.

[64] Tur, J., Pereira-Lopes, S., Vico, T., Marín, E.A., Muñoz, J.P., Hernández-Alvarez, M., et al., 2020. Mitofusin 2 in Macrophages Links Mitochondrial ROS Production, Cytokine Release, Phagocytosis, Autophagy, and Bactericidal Activity. Cell Reports 32(8): 108079.

[65] Kawalec, M., Boratyńska-Jasińska, A., Beresewicz, M., Dymkowska, D., Zabłocki, K., Zabłocka, B., 2015. Mitofusin 2 deficiency affects energy metabolism and mitochondrial biogenesis in MEF cells. PloS One 10(7): 1–18.

[66] Robb, E.L., Hall, A.R., Prime, T.A., Eaton, S., Szibor, M., Viscomi, C., et al., 2018. Control of mitochondrial superoxide production by reverse electron transport at complex I. The Journal of Biological Chemistry 293(25): 9869–79.

[67] Pryde, K.R., Hirst, J., 2011. Superoxide is produced by the reduced flavin in mitochondrial complex I: a single, unified mechanism that applies during both forward and reverse electron transfer. The Journal of Biological Chemistry 286(20): 18056–65.

[68] Scherer, P.E., Williams, S., Fogliano, M., Baldini, G., Lodish, H.F., 1995. A novel serum protein similar to C1q, produced exclusively in adipocytes. The Journal of Biological Chemistry 270(45): 26746–9.

[69] Karczewski, K.J., Francioli, L.C., Tiao, G., Cummings, B.B., Alföldi, J., Wang, Q., et al., 2020. The mutational constraint spectrum quantified from variation in 141,456 humans. Nature 581(7809): 434– 43.

